# Cell-type specific impact of glucocorticoid receptor activation on the developing brain

**DOI:** 10.1101/2020.01.09.897868

**Authors:** Cristiana Cruceanu, Leander Dony, Anthi C. Krontira, David S. Fischer, Simone Roeh, Rossella Di Giaimo, Christina Kyrousi, Janine Arloth, Darina Czamara, Silvia Martinelli, Stefanie Wehner, Michael S. Breen, Maik Koedel, Susann Sauer, Monika Rex-Haffner, Silvia Cappello, Fabian J. Theis, Elisabeth B. Binder

## Abstract

A fine-tuned balance of glucocorticoid receptor (GR) activation is essential for organ formation, with disturbances influencing health outcomes. Excess GR-activation *in utero* has been linked to brain-related negative outcomes, with unclear underlying mechanisms, especially regarding cell-type specific effects. To address this, we used an *in vitro* model of fetal human brain, induced pluripotent-stem-cell-derived cerebral organoids, and mapped GR-activation effects using single-cell transcriptomics across development. Interestingly, neurons showed targeted regulation of differentiation- and maturation-related transcripts, suggesting a delay of these processes upon GR-activation. Uniquely in neurons, differentially-expressed transcripts were significantly enriched for genes associated with behavior-related phenotypes and disorders. This suggests that aberrant GR-activation could impact proper neuronal maturation, leading to increased disease susceptibility, through neurodevelopmental processes at the interface of genetic susceptibility and environmental exposure.

## Introduction

Human developmental trajectories are influenced by both genetic and environmental factors^1^. Understanding the molecular and cellular mechanisms of how these factors impact brain development has been difficult, given the restricted access to living human fetal brain tissue. An important environmental risk factor is over-activation of the glucocorticoid system during brain development, which has been related to several negative long-term health outcomes, including cognitive, behavioral and psychiatric outcomes^1^. The GC system mediates key processes in fetal organ development and glucocorticoid receptor (GR) activation by GCs like cortisol is essential for proper formation and maturation of all organs, especially the lung and brain^2^. GR functions as a transcription factor and upon activation translocates to the nucleus where it binds specific glucocorticoid response elements (GREs) that enhance or repress the transcriptional response of many genes in a cell-type-specific manner^3^.

Tightly-regulated GR-activation seems to be critical for normal development. This is supported by endogenous mechanisms responsible for limiting excess activity of maternal glucocorticoids in the fetus, including a pregnancy-induced rise in corticosteroid-binding globulin^4^ that limits free cortisol and the placental enzyme 11β-hydroxysteroid dehydrogenase 2 (11β-HSD2) which converts maternal glucocorticoids to inactive cortisone, thus limiting its activity in the developing fetus^5^. In addition, increased glucocorticoids exposure has been associated with detrimental outcomes. One example is synthetic glucocorticoid (sGC) use in pregnancy. When premature delivery is a risk for the developing fetus, current clinical guidelines recommend antenatal treatment with sGCs like betamethasone or dexamethasone^6^. sGCs are also given starting in the first trimester for fetuses at risk for congenital adrenal hyperplasia (CAH), which accounts for 1:15,000 births^7^. In recent years, antenatal sGC treatments have been on the rise, as 10% of pregnant women worldwide are at risk for preterm delivery, and most receive sGCs to promote fetal organ maturation^2,8^. There are clear benefits of glucocorticoid treatment on organ maturation and overall reduction of premature birth risk^9^, but increasingly more negative long-term outcomes have also been reported. sGCs readily cross the placenta^10^, thus potentially placing the fetus in danger of overexposure. Accumulating evidence from human cohorts indicates that fetal sGCs exposure in mid- to late pregnancy may result in adverse postnatal outcomes including hypertension^11^, cardiovascular disease^12^, dysregulated activity in the hypothalamic-pituitary-adrenal (HPA) axis and detrimental effects on brain structure and development^13^. Continued sGC treatment for CAH has also been linked to altered cognitive performance, including inattention, as well as increased fearfulness in childhood^7^. Furthermore, elevated endogenous maternal glucocorticoids have been linked to negative outcomes especially regarding brain development^14^. For example, elevated prenatal maternal cortisol was significantly associated with altered neonatal amygdala connectivity, associated with differences in sensory processing and integration, and subsequently internalizing symptoms in girls^15^. In addition, conditions associated with increased maternal GCs and decreased placental 11β-HSD2, such as prenatal stress and prenatal maternal depression^1^, which could lead to increased fetal GC exposure, have repeatedly been associated with long-term negative behavioral outcomes in children, including problems with attention and emotional regulation, as well as changes in brain structure and function, at birth and in childhood^16-18^. Prenatal depression alone affects 10% of women worldwide, with higher rates in developing countries^19^. Together with the increasing prevalence of prenatal sGCs administration, the societal impact of this problem is large and underlines the necessity for a better understanding of the mechanisms by which increased prenatal GR-activation in the developing brain leads to negative health outcomes. For this, model systems are necessary.

GR-activation can be modeled robustly *in vitro* and *in vivo* and is thus amenable to system-wide investigation. Rodent studies have demonstrated that sGCs exposure during gestation leads to brain development perturbations in GR-targeted brain regions – the medial prefrontal cortex, anterior cingulate cortex and hippocampus^20^ – as well as depression-like phenotypes and other negative outcomes in adult offspring^21^. While animal models have contributed important insight into the mechanisms of prenatal GR-activation, they have limited use for understanding some human-specific risk mechanisms. During brain development, important differences exist between primates and rodents, notably regarding the abundance, lineage complexity and proliferative potential of certain neural progenitor cell types. This translates at a macro scale into a lisencephalic brain in mice compared to the gyrification of the human brain and marked differences in cell-type distribution and self-renewal or differentiation patterns^22^. In addition, animal models do not allow mapping of human-specific GREs and their possible interaction with human genetic variation, given that such enhancers are poorly conserved across species. A second interesting model system consists of primary human neuronal progenitor cell lines^23,24^. While these allow investigation of human-specific genomic effects, they do not recapitulate the complex architecture and cell types seen in the multi-cellular human brain. Human model systems using induced pluripotent stem cells (iPSCs) represent promising tools for filling this gap. Specifically, iPSC-derived human cerebral organoids recapitulate three-dimensional complexity and cell-to-cell interactions^25,26^ in addition to a human genetic background, thus allowing to model environmental exposures in the complex and species-specific context of human brain development. Organoids recapitulate early-to-mid fetal brain development, with most similarities matching the first and early second trimester^27-29^. This period is known to be critical for neuron production, migration, connection, and differentiation^1^, but environmental impacts during these sensitive periods are not fully understood. In this study, we aimed to test whether human cerebral organoids could be model systems to study the impact of prenatal GC exposure on the developing brain. We established that pharmacological GR-activation in cerebral organoids leads to cell-type-specific transcriptional responses that map to GR-responsive genes and cell types, which may moderate risk for negative neurodevelopmental trajectories.

## Results

### I. Cerebral organoids model the early developing brain *in vitro* and express the molecular machinery for response to glucocorticoids

First, we confirmed that our cerebral organoids mapped onto brain development trajectories using bulk RNA sequencing (RNAseq) at seven successive developmental times from day 17 to day 158 (**Table S1; Supplementary Results**). We could delineate a distinct and robust developmental trajectory at the whole transcriptome level, with same-age samples clustering closest together and closer to samples of adjacent age; and with age-defining genes clearly clustering in a progressive temporal manner (**Figure 1A**).

**Figure 1:**
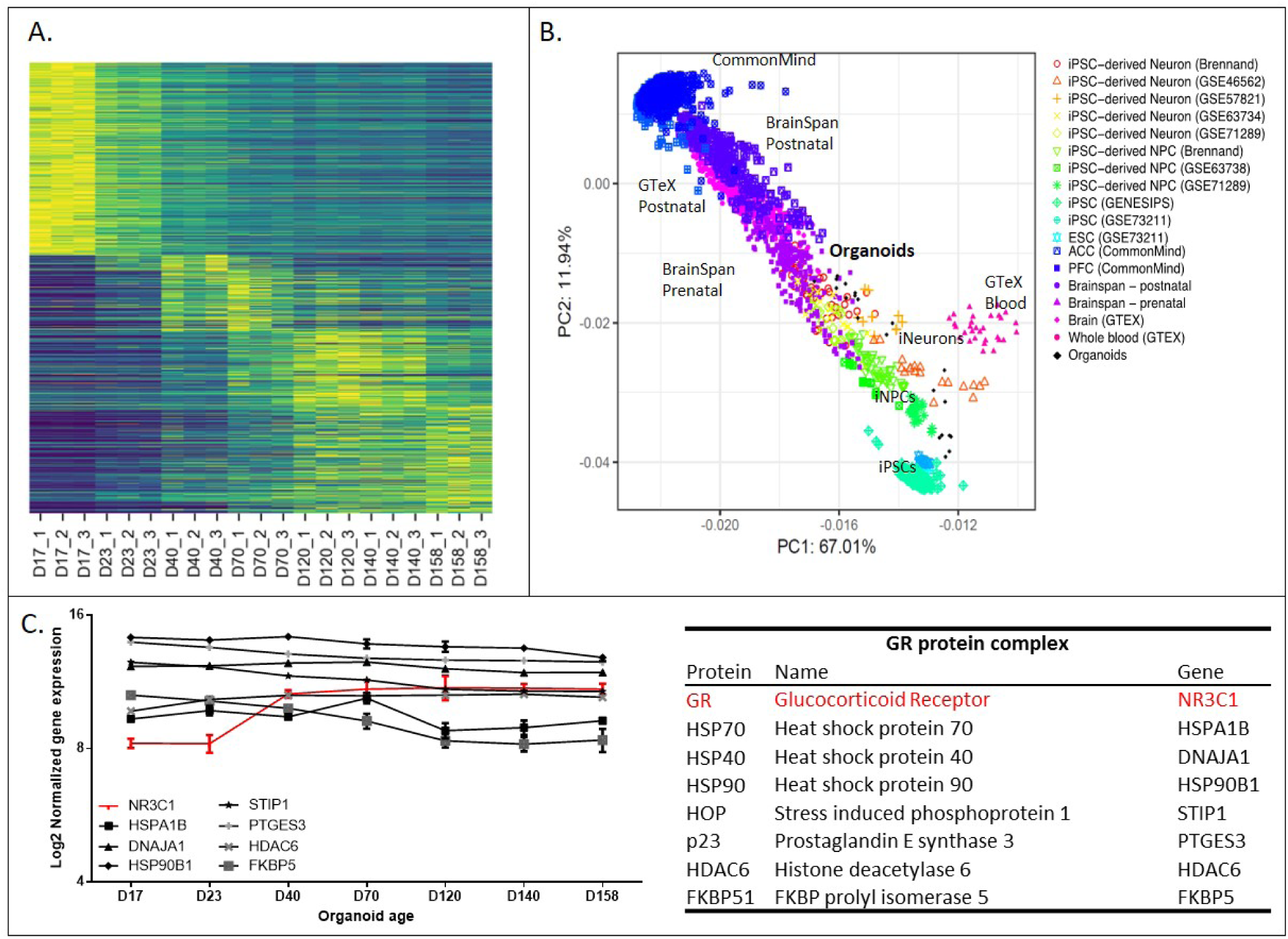
Transcriptional characterization of organoids’ developmental trajectory and GR-activation cellular machinery. **A.** Heatmap representing all genes peaking at each of seven tested time-points (D17-D158) of organoid culture (n=3), organized following temporal trajectory. Gene lists are in **Table S2**. Organoids show a robust and linear progression of gene expression across developmental time. **B.** Principle Component Analysis of published bulk transcriptome datasets highlighting mid-to-older age organoids positioned among fetal brain samples and advanced two-dimensional *in vitro* neuronal cultures. Organoids transcriptome datasets (n=21 total), represented by black diamond shapes, follow the linear trajectory of progressively maturing cell types or tissues according to organoid age. **C.** Expression levels of the primary GR protein complex genes across organoid development. Table (right) shows protein ID, gene name and gene ID. *NR3C1* expression significantly increased only between days 23 and 40 (ANOVA and Holm-Sidak post-hoc pair-wise test; p-value <0.0001; FC=1.29; n=3 per time point).

We compared these data with other bulk RNAseq datasets including fetal and adult postmortem brain^30,31^, as well as iPSC-derived *in vitro* differentiation models. Peripheral blood was used as a non-brain control. We found that older (beyond 70 days) organoids clustered with early fetal brain samples **(Figure 1B**). When surveying known markers cell-type associated with neuronal differentiation in the RNAseq data (**Figure S1A-B, Table S2** and **Supplemental Results**), as well as at the protein level using immunofluorescence staining (IF) (**Figure S1C**), we showed a gradual maturing over time of the three-dimensional brain-like structure centered around the functional unit of the ventricle.

We then asked whether the molecular machinery for GR-activation was present in this system, including essential chaperones and co-chaperones of the GR protein complex^32^ (**Figure 1C, Table S1**) using bulk RNAseq. We found *NR3C1* expression increased with organoid age, with a significant increase between days 23 and 40 (p-value<0.0001; FC=1.29), plateauing after day 40 (**Figure 1C**). Furthermore, the primary genes encoding the GR protein complex were abundantly and stably expressed across organoid development (**Figure 1C**), supporting the possibility for a functional GR complex in this system. Using IF, we also mapped the protein form of the *NR3C1* gene. We found GR protein expression throughout the organoids with no clear cell-type specificity (See **Figure 2** and next section).

**Figure 2:**
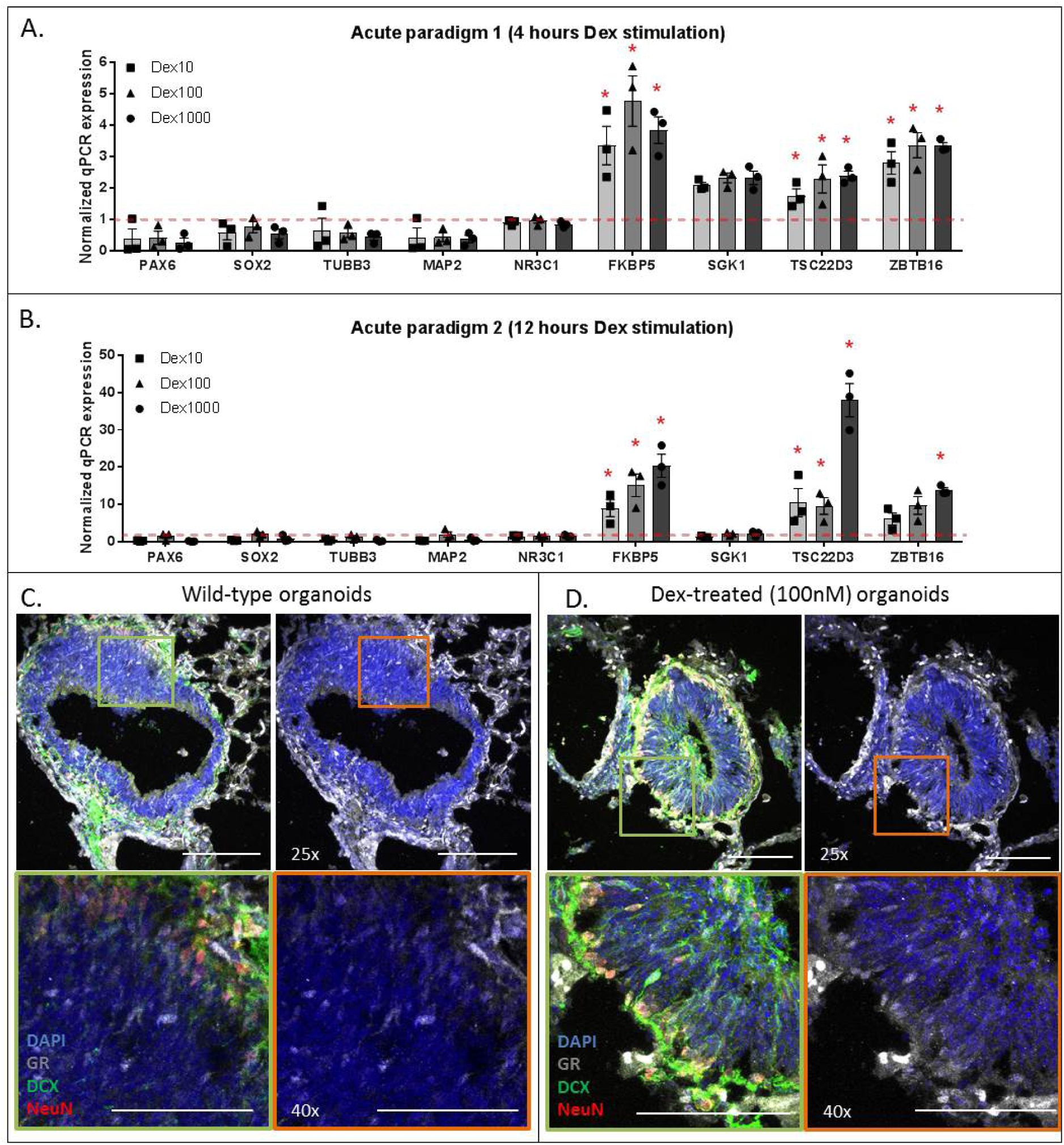
GR-activation paradigm in organoids. **A-B.** Quantitative PCR analysis of key GR-regulated genes across two different acute time paradigms 4 hours (panel **A**) and 12 hours (panel **B**) and dose (10nM, 100nM, and 1000nM Dexamethasone) stimulation paradigms. Organoids were stimulated at 45 days in culture (n=3). Gene expression was normalized to the geometric mean of endogenous genes GAPDH, POLR2A, and YWHAZ. Statistical analysis and results are reported in **Table S3**. Fold changes are reported in reference to Vehicle (DMSO). Expression is graphically represented was normalized to the Vehicle expression for each gene (indicated by the dotted red line). Statistical significance (two-sided) is denoted with * representing p-value ≤ 0.05. Error bars represent mean±SEM. **C.** Immunofluorescence image of an organoid in wild-type conditions showing GR protein localization in the cytoplasm of a neuronal cell (nuclei-DAPI = blue; DCX-neurons = green; GR = white). **D.** Immunofluorescence image of an organoid following 100nM Dexamethasone treatment for 12 hours showing nuclear translocation of GR in a neuronal cell (nuclei-DAPI = blue; DCX-neurons = green; GR = white; Magnification: upper panels = 25x, lower panels = 40x).

### II. Acute glucocorticoid exposure in cerebral organoids elicits differential expression of established glucocorticoid-responsive genes

We activated the GR in this *in vitro* model via exposure to the selective synthetic agonist dexamethasone (Dex). Organoids were grown for 45 days to ensure maximal GR expression, and two acute paradigms (4 and 12 hours) were tested, with three different Dex concentrations (10 nM, 100 nM, and 1,000 nM) in triplicate compared to vehicle (DMSO). Quantitative PCR analyses revealed that neither progenitor (*SOX2, PAX6*) nor neuronal cell markers (*TUBB3, MAP2*), nor the GR itself (*NR3C1*) were significantly Dex-reactive (**Figure 2A-B**; statistics in **Table S3**), suggesting that acute Dex treatment does not impact cell-type composition. This was contrary to transcripts known to be up-regulated by glucocorticoids: *FKBP5*^*33*^, *SGK1*^*22*^, *TSC22D3*^*32*^ and *ZBTB16*^*34*^. Most of the dose-time combinations tested resulted in significant up-regulation of these genes, except for *SGK1* (**Figure 2A-B**; statistics in **Table S3**). Twelve hours of stimulation (**Figure 2B)** elicited the strongest response in organoids, with fold-changes (Dex/Vehicle) as high as 39-fold for *TSC22D3* and 20-fold for *FKBP5* (**Table S3**). From these data, we concluded that 100 nM Dex for 12 hours represented a suitable acute GR-activation paradigm in cerebral organoids. Following this exposure, we also found evidence for GR-activation^31^ at the protein level. In the vehicle condition, GRs were localized in both the cytoplasm and nucleus (**Figure 2C**), while following Dex (100nM; 12h) all detectable GRs were in the nucleus (**Figure 2D)**, suggesting Dex-induced nuclear translocation of this transcription factor.

### III. Cell-type distribution in cerebral organoids across developmental time shows neuronal maturation

We next wanted to understand cell-type-specific glucocorticoids-responses in the organoid model. For this we performed single-cell transcriptome sequencing at three time-points that captured the developmental pattern comparable with fetal brain and GR expression trajectory in organoids; at 30, 60 or 90 days in culture (D30, D60, D90; n=4) in organoids exposed to vehicle (DMSO; Veh) or Dex (100 nM; 12 hours). We profiled over 15,000 cells, and following dissociation, microfluidic separation, selection of single-cells and quality control of sequencing data, we were left with 14,002 cells across all time-points and treatment conditions (D30 n=5,035 cells; D60 n=4,315 cells; D90 n=4,652 cells).

We applied graph-based cluster analysis using Scanpy^35^ on all 14,002 cells and clustered the dataset into 17 groups to discretize transcriptomic variability (**Figure 3A**). Of these, 13 clusters could be mapped to specific cell-types based on top differentially-expressed cluster markers as well as established protein makers consistent with recent publications (**Figure 3B-C; Table S4; Figure S2**)^25,26,36^. When separating cells by developmental time-point, by-and-large the same cell-types were represented and identified by the same markers, with the main difference being relative cell abundance (**Figure 3C; Figure S3**). For example, the proportion of cycling Neuroepithelial cells (cNE) decreased as the organoids aged (Chi-square=15.58, p-value=0.0004) while the proportion of more differentiated cell-types like the Dorsal Neurons (DN) increased (Chi-square=6.03, p-value=0.049) (**Figure 3C**).

**Figure 3:**
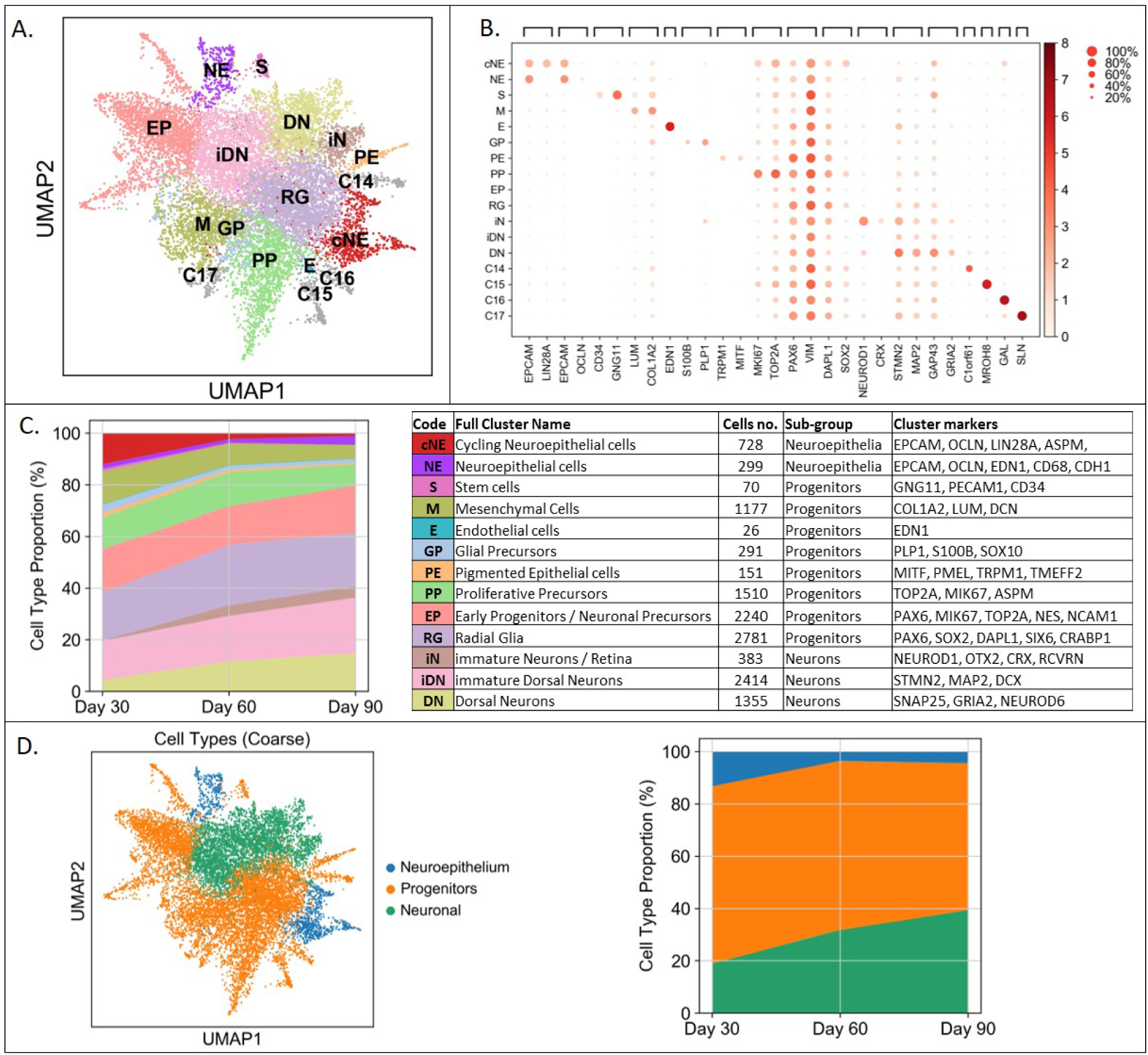
Cell-type characterization using single-cell RNA sequencing. **A.** Dimensionality reduction Uniform Manifold Approximation and Projection (UMAP) plot depicting 14002 single cells passing quality control from all time-points and treatment conditions (n=12 experiments). Colors depict each of 17 Louvain groups representing individual cell types and labeled with an abbreviation as defined in **C**, inset table. **B.** Dot plot showing the average expression by cluster and percent of cells expressing each of the top 1-2 genes defining each cluster. **C.** Left, cell-cluster distribution across organoid age (day 30, 60, 90). Right, definition of the 14 identifiable cell types, abbreviations, number of cells in each cluster, cell class assigned to, and the genes defining each cluster identity. **D.** Left, classification of cells into three cell classes (Neuroepithelial cells, Progenitors, and Neurons). Right, distribution of cell classes across organoid age (day 30, 60, 90).

All 17 cell clusters could be grouped into three global cell classes based on well-established markers as described in detail in **Supplemental Results**. These were Neuroepithelial cells (n=1,027), Progenitors (n=8,823) and Neurons (n=4,152) (**Figure 3D**). Even more prominently than with individual cell-types, the percent of cells in each cell class across organoid age changed with maturity. Neuroepithelial cells decreased from 13% at D30 to 4% cells D60 or D90 (Chi-square=8.3, p-value=0.016) while there was not significant change in Progenitors. Neurons increased progressively with organoid age from 19% to 32% to 39% of cells at D30, D60, and D90, respectively (Chi-square=9.8, p-value=0.007; **Figure 3D**).

### IV. Cell-type specific responses to acute glucocorticoids stimulation reflect a delay in neuronal maturation

We next evaluated cell-type-specific GR-activation responses using MAST^37^ for differential expression (DE) analysis. Given the comparable cell-types observed at D30, D60 and D90 (**Figure S3, Tables S5-S7**) we performed the main DE analyses in all combined cells to maximize statistical power, including technical batch as a covariate. We found no cell types unique to any treatment condition (**Figure 4A**). The GR (*NR3C1* gene, **Figure 4B**) was expressed across cell-types (see also **Figure S3** for time-point-specific results), though on average GR was more abundant in the NE, S, and M clusters, with over 50% of cells having detectable expression (**Figure 4C**). At the 17-cluster level, we found significant DE genes in 9 clusters, ranging from one to 55 DE transcripts (q-value≤0.05; **Figure 4D, Table S8**). The number of DE transcripts was not correlated with average *NR3C1* expression in the clusters (R^2^=0.001). However, absolute fold-changes for DE transcripts were significantly larger in *NR3C1*-positive cells within each of the three cell classes (**Figure S4**). Radial Glia (RG) had the most significantly DE genes (n=55) followed by the immature Dorsal Neurons (iDN, n=47 genes), the Dorsal Neurons (DN, n=38 genes), and the Proliferative Progenitors (PP, n=37 genes) (q-value≤0.05; **Figure 4D, Table S8**). No gene was DE in more than four clusters, but 5 genes were shared among the four clusters with most DE transcripts (PP, RG, iDN, and DN) (**Figure 4E**). Of these, one transcript (*MGARP*) was up-regulated, while 4 transcripts (*C1orf61, LINC01551, NEUROD6, NFIA*) were down-regulated by Dex. The direction of DE fold-change (FC) was consistent across all three time-points (**Tables S5-7**), indicating a cell-type and not temporally-dependent effect.

**Figure 4:**
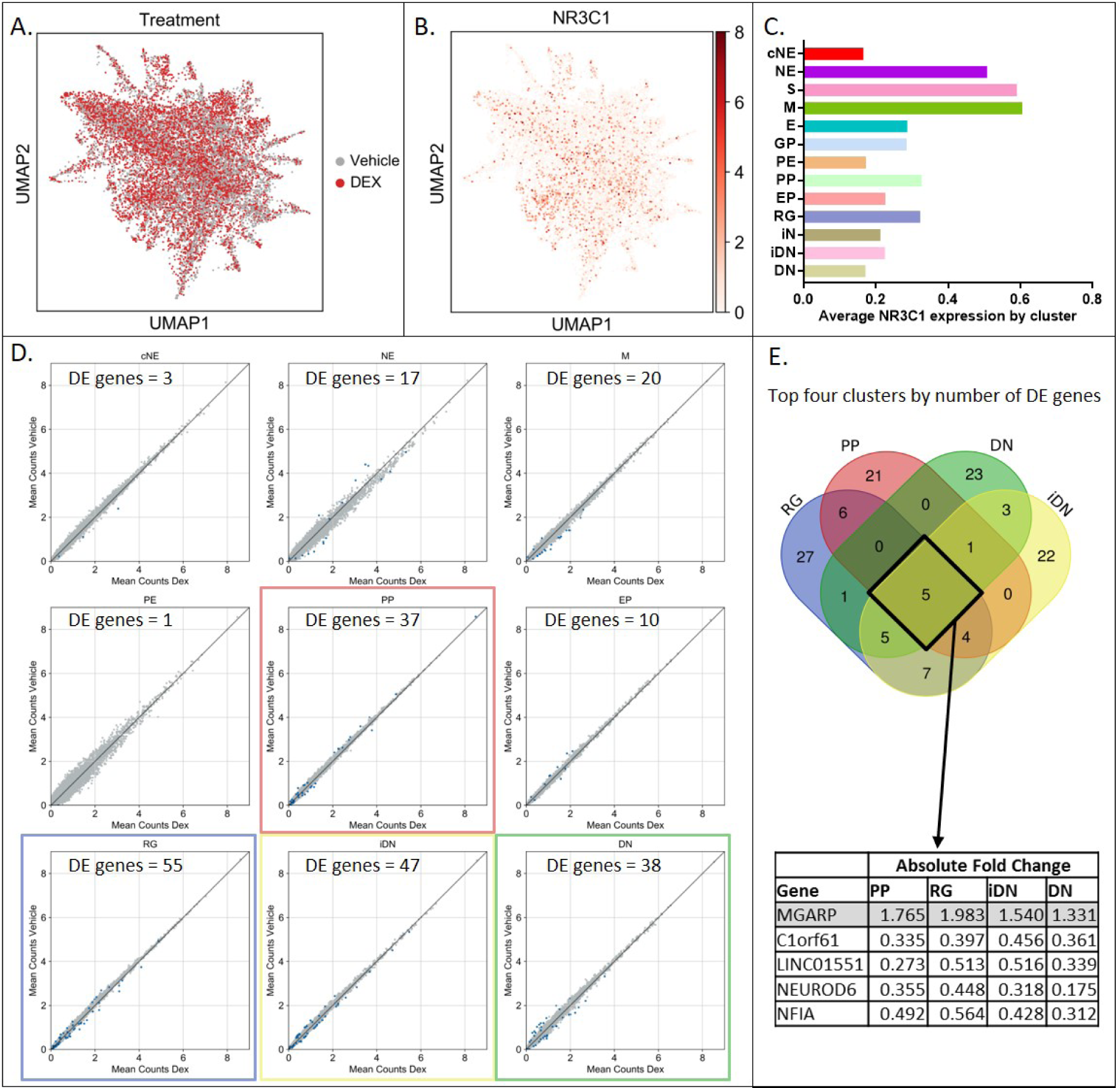
Cell-type-specific differential gene expression in organoids. **A.** UMAP plot depicting the cells from organoids that received treatment (red, n=6800 cells) or vehicle (grey, n=7202 cells). Even distribution of treated/untreated cells. **B.** UMAP plot depicting expression of the GR gene *NR3C1* across all 14,002 cells in the analysis. Even distribution of *NR3C1*-positive cells across cell types. **C.** Average *NR3C1* expression by cell cluster. Y-axis is cell-type abbreviation, x-axis is log2-normalized average gene expression across each cluster. Color scheme corresponds to Figure 4. **D.** Differential gene expression by cluster. Scatter plots (log2 fold change) of all genes differentially expressed between treatment (Dex, x-axis) and control (Veh, y-axis) where the significant genes (q-value ≤ 0.05) are shown by blue dots (n of DE genes marked on graphs)**. E.** Venn diagram depicting the intersection between lists of DE genes for the top four clusters by number of DE genes (PP, RG, iDN, and DN). The 5 genes shared by all four clusters are highlighted in the table along their differential expression fold change for each cluster.

When analyzing DE transcripts within the three broader cell classes, we found 68 DE genes in Neuroepithelial cells, 1,237 DE genes in Progenitors and 322 DE genes in Neurons (q-value≤0.05; **Figure 5A; Table S8)**. Only 14 DE genes were shared among all three cell classes, while 34 genes overlapped between Neuroepithelial cells and Progenitors, and 137 genes between Progenitors and Neurons (**Figure S6**). Most overlapping genes showed the same direction of regulation between cell classes, but 5 genes had opposite directions between Neurons and Progenitors. These were *CRABP1, DAPL1, ENO2, HES6*, and *PAX6* (**Figure 5B**), all of which are important in neurodevelopment, either by regulating cell proliferation or maintaining the progenitor pool^38-42^. Interestingly, all 5 genes were up-regulated in Neurons and down-regulated in Progenitors. When breaking down this effect by individual time-point (D30, D60, D90), all genes were consistently down-regulated (significant when q-value≤0.05) in Neurons, while in Progenitors this was only true for ENO2 and HES6. Together with an enrichment of common GO terms in DE transcripts from both Progenitors and Neurons which included cell differentiation, head development, neurogenesis, neuron differentiation and nervous system development (**Table S9, Supplementary Results**), these findings suggest that sustained GR-activation during neurodevelopment may interfere with neuronal differentiation and maturation in the long term, confirming previous data from 2-dimensional cell systems^24^.

**Figure 5:**
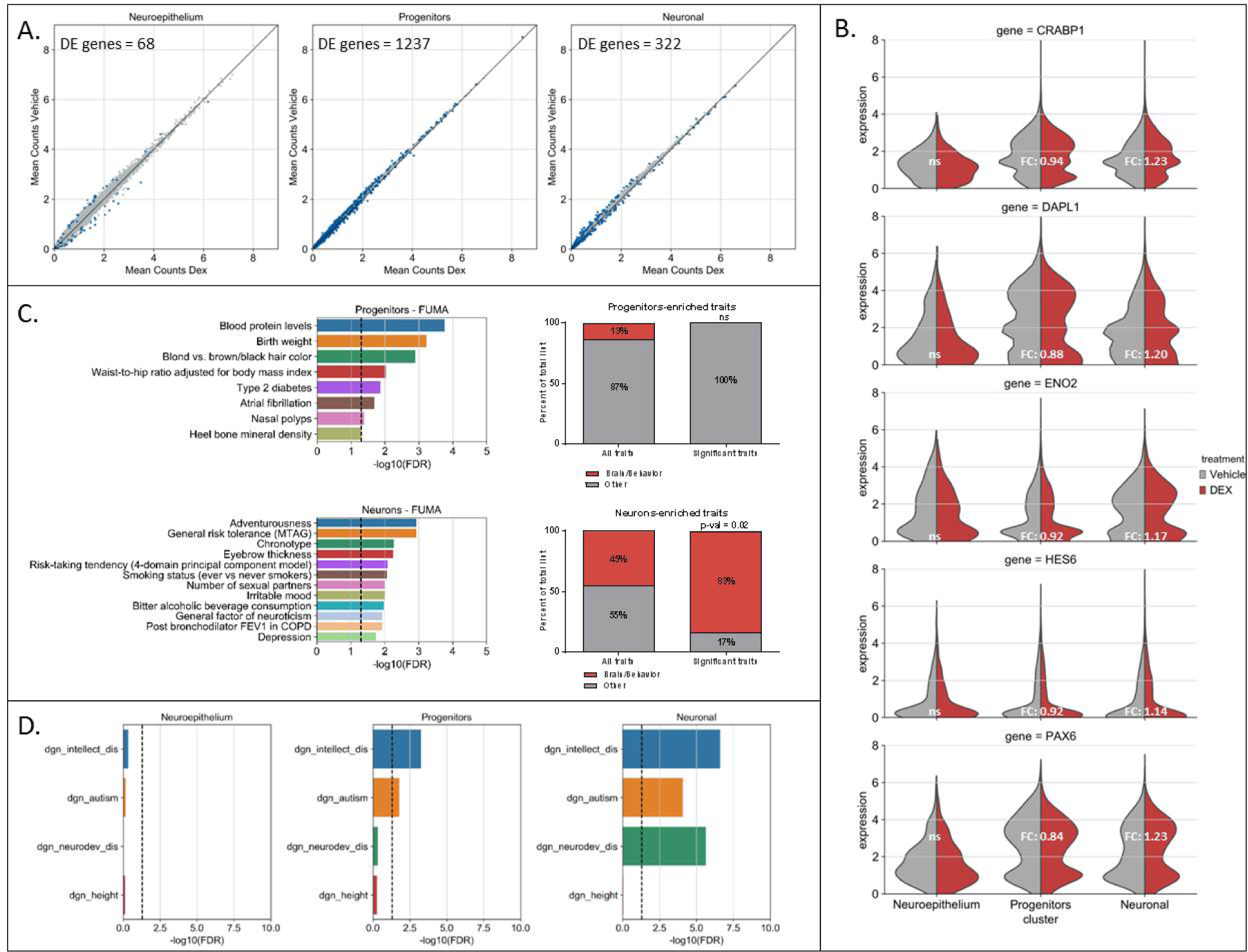
Phenotypic interpretation of differential GR response. **A.** Differential gene expression by cell class. Scatter plots (log2 fold change) of all genes differentially expressed between treatment (Dex, x-axis) and control (Veh, y-axis) where the significant genes (q-value ≤ 0.05) are shown by blue dots dots (n of DE genes marked on graphs). **B.** Split violin plots of the 5 genes that are significantly differentially expressed with fold changes in opposite directions between Progenitors and Neurons. **C.** Enrichment analysis in GWAS traits. Left, significant results from enrichment analysis of DE genes from Progenitors and Neurons versus all currently published traits with GWAS-significant results (q-value ≤ 0.05; Progenitors: n=8 significant traits; Neurons: n=12 significant traits). Right, comparison of significantly enriched traits classified as either ‘Brain/Behavior’ or ‘Other’ (% of total) in Progenitors or Neurons compared to the full traits lists with at least one gene overlap to Neurons (n=40) or Progenitors (n=272) DE genes. **D.** Enrichment analysis of neurodevelopmental disorders. Significant results from enrichment analysis of DE genes from Neuroepithelial cells, Progenitors and Neurons versus gene lists from the DisGeNET database for Intellectual Disability (ID), Autism Spectrum Disorders (ASD) and Neurodevelopmental Disorders (ND), as well as Height as a control. Dotted line represents the FDR-corrected significance threshold (q-value ≤ 0.05).

### V. Transcripts regulated by Dex in Neurons are specifically enriched for genes implicated by GWAS for behavioral traits

To better gauge a potential impact of aberrant GR-activation during neurodevelopment on phenotypes later in life, we mapped transcripts differentially regulated by Dex in our model of the developing brain to genes implicated by over 700 genome-wide association studies (GWAS) in different quantitative traits and diseases recorded in the GWAS Catalog^43^ (**Table S10**). We tested enrichment of our DE gene lists from the three cell classes (cut-off q-value≤0.05) within genes significantly associated with GWAS traits using FUMA^44^. No significant enrichment emerged for genes DE in Neuroepithelial cells, while transcripts DE in Progenitors were enriched for genome-wide associated genes in 8 traits and Neuron DE transcripts for genes associated with 12 traits (enrichment q-value≤0.05; **Table S10, Figure 5C**). We classified GWAS phenotypes not related to brain function as ‘Other’, while brain function-related traits were labeled ‘Brain/Behavior’. The latter were over-represented in traits enriched for Neuron DE transcripts, making up 83% of the traits with significant enrichment for this cell class (**Figure 5C** right; **Table S10**). This distribution was significantly different from the full list of traits (n=40) with at least one gene overlap to Neurons DE genes (Chi-square 5.46, p-value=0.02; **Figure 5C**) used as background for this analysis. Traits classified as ‘Brain/Behavior’ included depression, neuroticism, chronotype and adventurousness, suggesting that GR activation in the developing brain can impact genes relevant for behavioral phenotypes and psychiatric disorders later in life. Such an effect was not found in Progenitors, where all significant traits fell in the ‘Other’ category (**Figure 5C** right; **Table S10**).

Since prenatal GR exposure has been shown to increase risk for a number of psychiatric traits and disorders, we tested the enrichment of our DE gene lists among genes carrying common variants identified by the genome-wide meta-analysis of the Cross-Disorder Group of the Psychiatric Genomics Consortium, which included cases with 8 mental illnesses^45^ (gene list in **Table S11**). In this analysis, we found a significant enrichment only for DE genes in Neurons (q-value=0.0047 and OR=3.91; **Table S11**). To test whether the enrichment results could be reflective of cell-type-specific mean expression ranges in each cell class, rather than being linked to GR-activation we used a permutations-based approach, explained in **Online Methods** In this analysis gene sets comparable in size and mean expression distribution, but not significantly DE (cut-off q-value≤0.05) following Dex treatment were tested. A significant enrichment of these gene sets was only observed in 4 of 1,000 iterations (corrected empirical p-value=0.015; **Table S11**). This indicates that the enrichment of DE genes among genes with genome-wide significant associations in the Cross-Disorders GWAS is specific to the GR-responsiveness of the transcripts and not their expression level in Neurons.

### VI. GR-activation in Neurons specifically regulates neurodevelopmental disease-related genes

Neurodevelopmental phenotypes are not well represented among the tested GWAS traits^43^. To understand the relationship between our GR-activation findings in organoids and neurodevelopmentally-relevant genes, we tested for enrichment among genes associated with neurodevelopmental disorders. Firstly, we used gene lists based on various biological evidence from the DisGeNET database^46^, associated with three phenotypes: Intellectual Disability (ID), Autism Spectrum Disorders (ASD) and Neurodevelopmental Disorders (ND), and Height as a non-brain-related control (gene lists in **Table S12**). Secondly, we selected genes carrying loss-of-function mutations associated with ID from the Developmental Brain Disorders Database (DBDD)^47^ (gene list in **Table S13**). We found a significant enrichment for ID and ASD genes in DE transcripts from Progenitors (ID: q-value=0.00057, OR=1.36; ASD: q-value=0.0165, OR=1.53; **Figure 5D, Table S12**). An even more pronounced result emerged for DE transcripts in Neurons, where we found a strong significant enrichment with all three neurodevelopmental phenotypes (ID: q-value=2.49×10^−7^, OR=2.25; ASD: 8.29×10^−5^, OR=2.96; ND: 2.37×10^−6^, OR=5.53; **Figure 5D, Table S12**). While size and expression-level-matched gene sets (see above) for Progenitors also showed significant enrichments, this was not true for Neuron gene sets. Here permuted enrichments for any of the three neurodevelopmental phenotypes never reached p-value≤ the nominal p-value, making this finding specific to the Dex-regulated genes in Neurons (empirical q-value=0.012; **Table S12**).

When testing for enrichments within genes with ID-related loss-of-function mutations in DBDD^47^, we found a highly significant enrichment of DE transcripts in Neurons only (q-value=2.05×10^−7^; OR=5.60 **Table S13**), finding which proved to be driven by GR-activation, with 0 of 1,000 permuted iterations of the test gene list reaching p-value≤ the nominal p-value (**Table S13**). This enrichment was based on 17 DE transcripts mapping to ID-associated genes with loss-of-function mutations, including the *NRXN1, TCF4, TBR1* and *PAX6* genes where multiple pathogenic mutations have been linked to neurodevelopmental disorders. When breaking the analysis down by cell-type, this result was consistent and even more pronounced in the most mature cell cluster, the Dorsal Neurons (q-value=5.05×10^−8^, OR=25.57; permutation-based empirical q-value=0.004; **Table S13**).

The fact that neuronal transcripts DE following GR-activation were significantly enriched among genes carrying common variants associated with behavioral phenotypes as well as rare variants associated with ID, suggests a possible convergence of environmental and genetic factors on the same neurodevelopmental pathways.

## Discussion

In epidemiological and clinical studies, GR-activation by prenatal maternal stress or the administration of sGCs during critical periods of human brain development has been associated with several negative health outcomes including endocrine dysfunction, brain structural and functional alterations, and emotional, behavioral, cognitive or mental health problems^14,48,49^. In this manuscript, we explored whether iPSC-derived human brain organoids could contribute to a better mechanistic understanding of the detrimental effects of aberrant *in utero* GCs exposure. We found that in this system the GR was expressed and functional across all cell types, translocating to the nucleus with activation by the GR agonist Dex, and triggering a profound transcriptional response. Using single-cell RNA-sequencing (scRNAseq), we found that GR-activation transcriptional response was cell-type-specific, specifically pointing to interference with neuronal differentiation and maturation. Transcripts responsive to GR-activation in Neurons were significantly enriched for genes associated with behavioral traits in GWAS and implicated in neurodevelopmental disorders, including genes with loss-of-function mutations related to intellectual disability. This suggests a possible convergence of environmental (glucocorticoid exposure *in utero*) and genetic factors on the same cell-type-specific neurodevelopmental pathways. Thus, brain organoids open the possibility to interrogate human-specific genetic vulnerability to prenatal stress and glucocorticoid exposure.

Previous studies suggested that cerebral organoids recapitulate, to a degree, transcriptomic and epigenomic profiles of the developing human brain^25,26,28,29^, specifically matching the first and early second trimester. We confirmed that organoids match transcriptomic profiles of early fetal brains based on possibly the densest temporal characterization of this system thus far using bulk RNAseq. We found temporal gene expression patterns to be consistent with gradually maturing brain-like tissue. Importantly, both transcriptome analyses showed the developmental trajectory of our organoid model to be robust between biological replicates and across time in culture. scRNAseq revealed that organoids of different developmental ages consisted of the same brain-specific cell types, but with shifting relative distribution across maturation.

After demonstrating the presence of GR and its molecular partners in our system, we showed that GR-activation elicited a robust transcriptional response and nuclear translocation of the receptor. GR-expressing cells in a given cell class showed a significantly higher transcriptional response than comparable cells without detectable GR, supporting direct effects of GR-activation. Even though within a cell class GR-positive cells showed a stronger transcriptional response to GR-activation, the number of significantly regulated transcripts in a cell class was not directly correlated with the number of GR-positive cells, suggesting that GR-responsiveness is moderated by additional, cell-type-specific factors. These could include the distinct presence of GR-cofactors or specific epigenetic changes.

We found the most profound and impactful GR-activation response in neurons, with decreased expression of neuronal-specific genes like *NEUROD6, FOXG1, TBR1, MYT1L* and *NFIA*, but an increase of progenitor-specific genes like *PAX6, HES6, ENO2, CRABP1* and *MGARP* across several cell-types. This result could only be identified in a complex multi-cellular system like organoids, which contain progressively maturing cell-types from early neuroepithelia to dorsal neurons. While it confirms some previous results from cell and animal models showing glucocorticoids’ involvement in neurogenesis by increasing proliferation while decreasing differentiation^23,24,50^, our study is the first to support that aberrant GR-activation could lead to neuronal maturation and differentiation delay in a complex human developing brain model.

Finally, we also explored whether glucocorticoids-responsive transcripts would mediate neurodevelopmental, behavioral and psychiatric phenotypes. We found this to be true for both common and rare genetic disease associations. We found an enrichment of GWAS genes linked to behavioral phenotypes, including psychiatric disorders, specific to transcripts DE following GR-activation in Neurons. In addition, we observed a significant enrichment of GR-activated transcripts among genes related to neurodevelopmental disorders including ID and ASD, again uniquely in Neurons. The fact that enrichment results for both genes associated with adult-onset behavioral disorders and neurodevelopmental disorders were specific to Neurons and not more immature cell-types suggests that alterations of *in utero* GR-activation could lead to negative mental health outcomes mainly via effects in more mature cell-types – specifically by delaying maturation given our finding that many genes relevant for this process were down-regulated by GR-activation.

These results further highlight the interplay between environmental exposure like synthetic glucocorticoids or increased maternal cortisol and inherited genetic factors on nervous system development. Organoids as a model system allow exploration of the interaction between human-specific genetic variation and environmental exposures like prenatal maternal stress in a cerebral cell-type-specific manner. While this manuscript shows human cerebral organoids as suitable models for aspects of *in utero* glucocorticoid exposure, experiments in additional genetic backgrounds will be necessary for a more complete picture. Furthermore, future studies should explore effects of the endogenous ligand cortisol and other sGCs. In the absence of quantitative data from exposed fetal brain, it is currently not clear what concentration of sGC or endogenous maternal glucocorticoids reach the fetal brain in different circumstances, and this is certainly an important research area.

Our findings position cerebral organoids as important model systems to fulfill the research gap of how *in utero* environmental disturbances affect neurodevelopment leading to negative outcomes. Organoids also allow extending these investigations to genomics or cellular to functional investigations using electrophysiology. Finally, these systems allow the modeling of environmental exposure in the context of human-specific genetic variation. Organoids could thus contribute to improved intervention and prevention strategies for developmentally-determined neuropathology.

## Author Contributions

All authors contributed significantly to the data generation and preparation of this manuscript.

## Acknowledgements

This work was completed in part with support by a NARSAD Distinguished Investigator Grant to EBB, as well as an Alexander von Humboldt Foundation fellowship and a Banting Postdoctoral Fellowship to CC. We thank Dr. Barbara Treutlein and Sabina Kanton for their help with establishing the organoids single-cell isolation methods.

## Methods

### I. iPSCs and Organoids

Induced pluripotent stem cells were derived via reprogramming from NuFF3-RQ human newborn foreskin feeder fibroblasts (GSC-3404, GlobalStem)^51^. They were cultured on 6-well plates (Thermo Fisher) coated with 1:30 Matrigel (Corning) in mTesR1 basic medium supplemented with 1x mTesR1 supplement (Stem Cell Technologies) at 37°C, with 5% CO_2_. Cerebral organoids (COs) were generated as previously described^52^, starting with 9,000 iPSCs dissociated into single cells using StemPro Accutase Cell Dissociation Reagent (Life Technologies) per each well in U-shaped low attachment 96-well tissue culture plates (Corning) in hES medium (DMEM/F12-GlutaMAX supplemented with 20% Knockout Serum Replacement, 3% FBS, 1% Non-essential amino acids, 0.1 mM 2-mercaptoethanol, 4 ng/ml bFGF and 50 µM Rock inhibitor Y27632) for 6 days in order to form embryoid bodies (EBs). On day 6, EBs were transferred into low attachment 24-well plates in Neural Induction (NIM) medium (DMEM/F12 GlutaMAX-supplemented with 1:100 N2 supplement, 1% Non-essential amino acids and 5 µg/ml Heparin) and cultured for an additional 6 days. On day 12 EBs were embedded in Matrigel (Corning, 354234) drops and transferred to 10-cm tissue culture plates in Neural Differentiation medium (NDM) without Vitamin A medium (DMEM/F12GlutaMAX and Neurobasal in ratio 1:1 supplemented with 1:100 N2 supplement 1:100 B27 without Vitamin A, 0.5% Non-essential amino acids, insulin 2.5 µg/ml, 1:100 Antibiotic-Antimycotic and 50 µM 2-mercaptoethanol) in order to form organoids. 4 days after Matrigel embedding, organoids were transferred into an orbital shaker and cultured in NDM with Vitamin A (DMEM/F12GlutaMAX and Neurobasal in ratio 1:1 supplemented with 1:100 N2 supplement 1:100 B27 with Vitamin A, 0.5% Non-essential amino acids, insulin 2.5 µg/ml, 1:100 Antibiotic-Antimycotic and 50 µM 2-mercaptoethanol). Organoids were grown in these conditions at 37°C with 5% CO_2_ until collection for RNA extractions, cryopreservation, or single-cell dissociation.

### II. Glucocorticoid stimulation

Organoids of different ages depending on experiment, were treated with glucocorticoids (GCs) by dissolving Dexamethasone (Dex) to the appropriate concentration (10 nM, 100 nM, or 1000 nM) in DMSO and subsequently in the culture medium. For acute stimulation, 4 hours or 12 hours exposure time was employed. For chronic treatment, exposure lasted for 7 days with media changes every second day. All exposures were performed in triplicate experiments.

### III. Immunofluorescence

Organoids were fixed using 4% paraformaldehyde for 45 minutes at 4°C, cryopreserved with 30% sucrose, fixed in optimal cutting temperature (OCT) compound (Thermo Fisher Scientific) and stored at −20°C prior to cutting 16 um cryosections. For immunofluorescence, sections were post-fixed using 4% PFA for 10 mins and permeabilized with 0.3% Triton for 5 mins. Sections were subsequently blocked with 0.1% TWEEN, 10% Normal Goat Serum and 3% BSA. Primary and secondary antibodies were diluted in blocking solution, and fluorescent staining was visualized and analyzed using a Leica laser-scanning microscope. For staining with PAX6 and SATB2 the slides were put through antigen retrieval before fixing with paraformaldehyde. More specifically, the slides were incubated with citric buffer (0.01 M, pH 6.0) for 1 min at 720 Watt and 10 mins at 120 Watt, left to cool down at room temperature for 20 mins and washed once with PBS.

| Antigen | Dilution | Vendor | Catalog # |
| --- | --- | --- | --- |
| DAPI | 1:1000 | Sigma Aldrich | D9542 |
| PAX6 | 1:500 | Biozol | BLD-901301 |
| SOX2 | 1:500 | Cell Signaling | 2748 |
| HOPX | 1:1000 | Santa Cruz | HPA030180 |
| CTIP2 | 1:500 | Abcam | ab18465 |
| SATB2 | 1:500 | Abcam | Ab51502 |
| TBR1 | 1:500 | Abcam | ab31940 |
| MAP2 | 1:500 | Sigma Aldrich | M4403 |
| TBR2 | 1:500 | Abcam | ab23345 |
| NESTIN | 1:500 | Santa Cruz | sc23927 |
| GR (D8H2) | 1:100 | Cell Signaling | 3660S |

### IV. Bulk RNA sequencing

Organoids (following 17, 23, 40, 70, 120, 140, or 158 days in culture) were collected for bulk RNA extractions using the RNeasy Mini extraction kit (74104, Qiagen) according to the manufacturer’s instructions. RNA quality and concentrations were measured on an Agilent 2100 Bioanalyzer (Agilent). Three replicates were analyzed per time point, every sample containing 1-3 pooled organoids. Sequencing libraries were prepared from a starting amount of 100ng total RNA using the NEBNext^®^ Ultra™ DNA Library Prep Kit for Illumina (E7370L, New England Biolabs) using ribosomal depletion as a selection method, and sequenced paired-end on an Illumina HiSeq4000 system at the Helmholtz Zentrum Core Facility (Munich, DE). 5 libraries were pooled per lane for a total coverage of on average 50 M reads/library. Raw reads were processed using FastQC^53^ and cutadapt^54^ and aligned using the STAR aligner^55^. The counts data were batch-corrected, normalized and analyzed and using the ImpulseDE2 framework^56^. For practical representation in the heatmap (**Figure 1**), all genes were normalized to 1. Individual genes were plotted using GraphPad Prism V6 using log2-transformed values and mean ± standard deviation.

To confirm the developmental specificity of our cerebral organoid data, we integrated bulk gene expression data with several transcriptional data sets from human post-mortem brain tissue and other iPSC-derived models. We used a recently described approach to integrate these data^57,58^. In brief, a total of 15 independent studies were analyzed covering 2,716 independent samples and 11,572 genes. These studies span a broad collection of developmentally specific gene expression, covering expression related to iPSCs, iPSC-derived NPCs, iPSC-derived neurons, bulk prenatal brain tissue (early, mid and late fetal stages) as well as bulk postnatal brain tissue (early, mid and late stages). All expression values were converted to log_2_ RPKM and collectively normalized using quantile normalization from the *limma* R package^59^. These data, along with our bulk cerebral organoid expression data were jointly analyzed and integrated using principal component analysis (PCA). The first two principal components were used to depict developmental trajectories.

### V. Quantitative Real-time Polymerase Chain Reaction (qRT-PCR)

Total RNA was extracted from organoids using the RNeasy Mini extraction kit (74104, Qiagen) according to the manufacturer’s instructions. Complementary DNA (cDNA) synthesis was performed using the Maxima H Minus Reverse Transcriptase (Thermo Fisher) along with oligo(dT)16 primers (Invitrogen) and random hexamers (IDT DNA) in a 1:1 ratio. Real-time PCR reactions were run in quadruplicate using PrimeTime qPCR Primer Assays (IDT DNA) and PrimeTime^®^ Gene Expression Master Mix (IDT DNA) on an ABI PRISM 7900HT Sequence Detection System (Applied Biosystems) or a LightCycler 480 Instrument II (Roche). Relative gene expression levels were quantified using the absolute quantification method, and using *GAPDH, POLR2A* and *YWHAZ* as endogenous genes. Statistical differences between groups were analyzed by Two-Way repeated measures ANOVA with Dunnet’s post-hoc multiple comparisons tests. Statistical significance was calculated, and graphs were plotted using GraphPad Prism 7. A p-value of ≤0.05 was considered statistically significant and marked as *, and ≤0.1 was considered suggestive of a trend for significance and marked as # in graphs.

| Gene ID | Gene name | PrimeTime Assay |
| --- | --- | --- |
| SOX2 | SRY (sex determining region Y)-box 2 | Hs.PT.58.237897.g |
| PAX6 | Paired box protein Pax-6 | Hs.PT.58.38519242 |
| MAP2 | Microtubule-associated protein 2 | Hs.PT.58.20103440 |
| TUBB3 | Class III β-tubulin | Hs.PT.58.20385221 |
| NR3C1 | Glucocorticoid Receptor | Hs.PT.58.27480377 |
| FKBP5 | FKBP prolyl isomerase 5; FK506 binding protein 5 | Hs.PT.58.20523859 |
| SGK1 | Serum and glucocorticoid-regulated kinase 1 | Hs.PT.58.19153459.gs |
| TSC22D3 | GILZ; glucocorticoid (GC)-induced leucine zipper | Hs.PT.58.4331913 |
| ZBTB16 | Zinc finger and BTB domain containing 16 | Hs.PT.58.605743 |
| GAPDH | Glyceraldehyde 3-phosphate dehydrogenase | Hs.PT.39a.22214836 |
| POLR2A | RNA Polymerase II Subunit A | Hs.PT.39a.19639531 |
| YWHAZ | Tyrosine 3-Monooxygenase/Tryptophan 5-Monooxygenase Activation Protein Zeta | Hs.PT.39a.22214858 |

### VI. Single-cell RNA sequencing

#### Data collection

Whole organoids (after 30, 60, or 90 days in culture) were treated with Dex (100 µM) or Vehicle (DMSO) for 12 hours until harvesting for single-cell preparation. All experiments were performed in quadruplicate at all three time points, with a paired design of a treatment (Dex) or control (Veh) condition per each experiment/iCell8 chip, for a total of 12 chips. Single cells were dissociated using StemPro Accutase Cell Dissociation Reagent (Life Technologies), filtered through 30 uM and 20 uM filters (Miltenyi Biotec) and cleaned of debris using a Percoll (Sigma, P1644) gradient. Single cells were resuspended in ice-cold Phosphate-Buffered Saline (PBS) supplemented with 0.04% Bovine Serum Albumin and prepared for single-cell separation and labeling using the iCELL8 Single-Cell System (Wafergen, Takara Bio) according to the manufacturer’s recommendations. Briefly, cells were stained with Propidium Iodide (for live cells) and DAPI (for dead cells) on ice for 5-10 minutes and dispensed in the loading plate. Following microfluidic separation, iCell8 chips were imaged using the built-in fluorescence microscope and snap-frozen and stored at −80°C until library preparation. Based on fluorescence labeling, only wells containing live single cells were selected for library preparation, which was performed according to the manufacturer’s guidelines using the iCell8 Chip and Reagent Kit using in-chip RT-PCR amplification chemistry (Wafergen, Takara Bio) and Nextera XT DNA Library Preparation Kit and Nextera XT Index Kit (Illumina). Libraries were assessed using a High Sensitivity DNA Analysis Kit for the 2100 Bioanalyzer (Agilent) and KAPA Library Quantification kit for Illumina (KAPA Biosystems), and sequencing was performed paired-end with 26nt/100nt configuration on an Illumina HiSeq4000 system at the Helmholtz Zentrum Sequencing Core Facility (Munich, DE).

## Data Analysis

### Pre-processing and quality control of single-cell RNAseq data

Sequencing of the iCell8 single cell libraries was performed on an Illumina HiSeq4000 (Illumina, San Diego, CA), 1 lane per chip, generating paired-end reads of 100 bp length. The initial quality check was performed using FastQC^53^ before demultiplexing the cells by their barcode with *Je* multiplexer version 1.0.6^60^ requiring a perfect match of the sequence. Subsequent adaptor trimming was performed using cutadapt version 1.11^61^. For read alignment, the STAR module^55^ of the Cell Ranger 2.1.1 release with the corresponding reference index of GRCh38 version 1.2.0 was used, both provided by the 10X Genomics. Only uniquely aligned reads with a minimum overlap of 31 bp to the transcriptome were regarded for gene quantification with the featureCounts package version 1.6.4^62^.

We carried out all downstream analyses using the python-based Scanpy package^35^ (https://github.com/theislab/scanpy) unless stated otherwise. We converted Ensemble Gene IDs to Gene Symbols using the R package BED^63^. To remove low quality cells, we filtered cells with a high fraction of reads from mitochondrial genes (20% or more - indicating stressed or dying cells), cells with <2,000 or > 600,000 total reads, as well as cells expressing < 600 genes. We also removed cells that were labelled as empty wells as part of the iCell8 library preparation procedure. In addition, we excluded genes with expression in < 20 cells. When analysing cells from one of the three time-points separately we removed all genes with expression in < 10 cells of the time-point in question. Unless stated otherwise, we carried out all following analyses with cells from all time-points combined as well as separately for each of the three time-points (day 30, 60 and 90).

### Normalisation and batch correction

To approximate the effect of sequencing depth in the data, we used the *computeSumFactors()* function from the R package scran^64^. To remove this effect, we divided the reads of each cell by its associated size factor as computed by scran. We log-transformed the data and used the resulting expression matrix for computing marker genes as well as for differential expression analysis downstream. For the visualisation of gene expression levels of individual genes we removed technical variation introduced by handling of the different iCell8 chips using the Scanpy implementation of the combat package^65^. For improved interpretability of plots representing gene expression, we limited colour scales and axis limits to combat-corrected expression values between 0 and 8. This ensures comparability between the visualisation of the expression values of different genes prevents outlier cells with extremely high expression values from dominating visualisations and also ensures that negative expression values produced by combat are not affecting visualisations.

The small positive and negative expression values introduced by combat in places of zero gene expression are particularly problematic when relying on dot plots to visualise the expression of marker genes. While this behaviour is a natural result of the underlying regression approach combat uses to remove technical variation, this result is not compatible with a dot plot as a marker gene visualisation tool. This is because the concept of a dot-plot relies heavily on the absence of expression of a gene in all but a few cells. In order to ensure compatibility between combat-correction and dot plot visualisations of marker genes, we adjusted the expression matrix in the following way: (1) we computed the median expression of each gene after combat correction while excluding all values from the median calculation that were zero before combat correction; (2) we set all entries in the post-combat expression matrix to zero which were zero before combat correction and at the same time were below the previously computed median expression value of the respective gene. This way, we only allowed combat to activate the expression of genes if the combat-corrected expression value was above the median expression value of that gene. On the one hand this prevents combat to turn on expression of spurious genes by a small amount and on the other hand still allows for turning on gene expression if the batch effect dominates the expression of a particular gene.

In order to obtain a meaningful low-dimensional embedding of the data, we carried out a separate normalisation and batch correction step starting again from the raw count data as before. We used a negative binomial generalised linear regression model with regularised over-dispersion parameter theta, as introduced in^66^, with the iCell8 chip ID as a batch covariate. To this end we used the R function *norm.nb.reg ()* from the github repository associated with^66^ (https://github.com/ChristophH/in-lineage). We constructed the normalised expression matrix from the Pearson residuals of the regression model.

### Low dimensional embedding, visualisation and clustering

We computed the single-cell neighbourhood graph on the 50 first principal components of the negative binomial residuals’ expression matrix using 15 nearest neighbours. We used Uniform Manifold Approximation and Projection (UMAP)^67^ for visualising the data in two dimensions. For clustering and cell type identification we used louvain-based clustering^68^ at varying resolution in different parts of the data manifold as implemented in louvain-igraph (https://github.com/vtraag/louvain-igraph) and adopted by Scanpy. We annotated cell types based on the expression of known marker genes. We merged clusters if only reflecting further heterogeneity within a cell type not discussed in this manuscript. For the exact steps of clustering and annotation and the parameters used consult the available code. In different figures varying resolutions of subtype clustering are shown in this manuscript. We identified characteristic gene signatures of each cluster by testing for differential expression of a single cluster against all other cells using a t-test with overestimated variance implemented in the sc.*tl.rank_genes_groups()* function of Scanpy.

### Differential Expression Testing

We used the MAST^69^ package for all single-cell differential expression analyses. MAST uses a two-part linear (‘hurdle’) model to match the distribution of scRNA sequencing data. We tested differential expression between Dex and Vehicle samples in different subgroups of cells. Firstly, we tested the effect in each of the fine clusters individually. Secondly, we investigated the same effect, but instead considering the three coarser cell classes. To correct for the batch effect of iCell8 individual experiments/chips, we included the chip ID as a covariate in the MAST model. Additionally, we included the number of genes expressed in each cell as a further covariate. After fitting the model we used a likelihood ratio test to test for differential expression between conditions. As an output from MAST we obtained raw p-values, p-values corrected for multiple testing with the Benjamini-Hochberg approach (false discovery rate) and log2 fold changes of gene expression between conditions. We used a false discovery rate of 0.05 as a significance threshold.

### Treatment response in NR3C1-positive vs. -negative cells

To investigate dependence of the Dex response on expression of the GR, we fitted the MAST model as described above for each of the three coarser cell classes and obtained log2foldchanges between cells that do or do not express the gene *NR3C1* (GR). As a next step, we intersected the full gene list with genes previously identified to be differentially responsive to Dex in hippocampal progenitors^24^. We then used the Wilcoxon signed-ranks test to ask if there was a significant difference in the response between NR3C1-positive and -negative cells in each of the three cell-type groups. Results were deemed significant at a p-value ≤ 0.05. This analysis was done for cells from all three time-points combined and not for individual time-points.

### Enrichment analyses

Enrichment analyses were carried out only on the differential expression results of cells from all three time-points combined, not on DE results from individual time-points.

#### a. FUMA

Enrichment of differentially-expressed genes against genes carrying SNPs with Genome-Wide Association to a variety of traits was tested using the FUMA algorythm^44^ by inputting the various DE gene lists into the GENE2FUNC software. This analysis references the NHGRI-EBI GWAS Catalogue^43^ (https://www.ebi.ac.uk/gwas/) most recently updated on 27 May 2019 and containing 943 traits from different studies. We ran an enrichment analysis separately for the DE genes (FDR ≤ 0.05) in the Neuroepithelia (genes recognized by software: Input: 66; Background: 16,752), Progenitors (genes recognized by software: Input: 1163; Background: 16837) and Neurons (genes recognized by software: Input: 303; Background: 16,829) cells classes and limited the analysis to traits where 5 or more genes overlapped with the respective test gene sets (corrected p ≤ 0.05). There were no traits with overlapping genes for the Neuroepithelial cells DE genes, so we continued the analyses with the other two gene sets. We categorized all traits as ‘brain/behaviour’ or ‘other’ in the full list of traits for each cell class where at least one gene was shared. We assessed significant divergence from the ‘master’ trait list using a Chi square test, with significance set at p ≤ 0.05.

#### b. Enrichment analyses for disease-associated genes

To test for enrichment of selected gene-sets in the differentially expressed genes (FDR ≤ 0.05), we used a hypergeometric test as implemented in the package diffxpy (https://github.com/theislab/diffxpy/). We tested for enrichment of all Gene Ontology Biological Function sets (v6.2), obtained from the Molecular Signatures Database^70,71^. We furthermore tested for enrichment of genes sets associated to disease as follows. **(1)** Genes mapping to SNPs with genome-wide significance in the genome-wide meta-analysis of the Cross-Disorder Group of the Psychiatric Genomics Consortium, which included 8 mental illnesses^45^ (gene list in **Table S11**). **(2)** Gene sets from the DisGeNET database^46^, which includes curated lists of disease-associated genes not only from genetic associations, but also from gene expression analyses and pharmacological studies. Here we focused on three phenotypes: Intellectual Disability (ID), Autism Spectrum Disorders (ASD) and Neurodevelopmental Disorders (ND), and used Height as a non-brain-related phenotype (Gene list in **Table S12. (3)** Genes carrying loss-of-function mutations and associated with Intellectual Disability from the most recent Developmental Brain Disorders Database (DBDB: https://www.dbdb.urmc.rochester.edu/home; Updated March 2019)^47^ (gene list in **Table S13**). We carried out enrichment tests only for cells from all time-points combined and in the following subgroupings: each of the three coarse cell classes individually or each of the fine clusters individually. We applied Benjamini-Hochberg FDR correction on the cluster level (correcting for multiple testing within different groups of cells) as well as on the enrichment list level for the number of compared gene lists when using more than one gene set at once as with sets obtained from the DisGeNET database.

#### c. Permutation testing of enrichment results

To ensure that enrichment analyses results were not simply reflective of high expression levels of marker genes in particular cell types or classes, we generated 1,000 permutations of equal-sized gene sets that also had the same mean expression distribution as the significant DE gene sets from each cell class (68 DE genes in Neuroepithelial cells, 1,237 DE genes in Progenitors and 322 genes in Neurons). The latter was achieved using 10 equal-gene-number bins where the mean gene expression was matched by bin. We then repeated all previous enrichment analyses using each of the permuted genes sets and counted the number of times a p-value ≤ the nominal p-value was reached. We used this to compute empirical p-values and corrected for the number of comparisons (both number of cell types/classes with significant enrichment results to consider, as well as the number of gene lists tested) using Bonferroni correction.

### Software specifications

We used Python v3.6.8 with Scanpy v1.4, anndata v0.6.18, h5py v2.9.0and diffxpy v0.6.3. Versions of packages required by Scanpy that might influence numerical results are indicated in the custom scripts. We used R version 3.6.0 with packages scran v1.12.0, MAST v1.10.0 and BED v1.1.5 with database version with UCB-Human 2019.04.23. We used matplotlib and seaborn to generate figures.

### Data and code

Primary data and processed data are available upon request prior to publication and will be made publicly available upon publication. Code for all analyses is available in the attached file.

## Supplementary Results and Figures

### Supplementary Results

#### Temporal expression of cell-type markers in bulk RNAseq data

We showed consistent levels of early progenitor markers PAX6 and NESTIN (*NES* gene) throughout organoid development. Intermediate progenitor markers TBR2 (*EOMES* gene) and HOPX progressively increased in expression across organoid maturation with significant peaks at day 158, likely reflective of increased neuronal maturation (**Figure S1A, Table S2**). Complementarily, young neuronal marker MAP2 and mature neuronal markers TBR1 and CTIP2 (*BCL11B* gene) appeared later in organoid development but maintained constant levels as the organoids increased in age, from day 40 on with a significant peak at day 120 (**Figure S1B, Table S2**). This cell-type distribution is consistent with a gradual maturing of the three-dimensional brain-like structure centered on the functional unit of the ventricle, with progenitors populating the apical surface of the ventricular zone and migrating toward the basal surface of the cortical plate as they differentiate. At the protein level following IF staining, this trend is clear based on early progenitor markers PAX6 and NESTIN, intermediate progenitor markers TBR2 and HOPX, early neuronal marker MAP2 and late neuronal markers TBR1 and CTIP2 at both 40 days and 90 days of organoids culture (**Figure S1C**).

#### Annotations of cell types based on marker genes in single-cell transcriptome data

All 17 identified cell clusters fit within three cell classes based on well-established markers^25,26,28,36,72-74^ as follows (**Figure 3B, Figure 3C** inset table). Neuroepithelia (n=1027 cells total), representing the most immature cells, were characterized by expression of epithelial marker *EPCAM* (**Figure S2**) and tight junction protein *OCLN*. Importantly, these early cells do not yet express vimentin (VIM) at high levels, which appears during the mesenchymal-to-epithelial transition from neuroepithelia to radial glia. These symmetrically-dividing cells represent the precursors to all potential neuronal and progenitor cell types, and form the neural plate *in vivo* and neural tube during embryonic development. In our organoids, we see two neuroepithelial clusters, with the cycling neuroepithelia (cNE) being primarily made up of day 30 cells and distinctly expressing cell cycling markers *LIN28A* and *ASPM*, while the slightly older neuroepithelia (NE) cluster distinctly expressing a combination of early markers *EDN1* (endothelial), *CD68* (associated to microglia and other peripheral cells), *TSTD1, CDH1*, all of which have been shown by other groups to be associated with NE cells (**Figure 3B; Figure S2**).

The second cell class represents the Progenitors (n=8823 total cells), defined by appearance of VIM^72^ (**Figure 3C-D; Figure S2**), and consisting of 12 clusters including most notably the Mesenchymal cells (M) which express structural proteins like *LUM* and *DCN* and collagens like *COL1A2*; Glial Precursors (GP) which express oligodendrocyte precursor as well as astrocyte marker *S100B* and glial precursor transcription factor *SOX10*; Proliferative Precursors (PP) defined by expression of cycling markers *TOP2A, MIK67, ASPM*; Early Progenitors / Neuronal Precursors (EP) which express VIM, progenitor marker PAX6 as well as neuronal lineage markers *NES, NCAM1*; and Radial Glia (RG) expressing markers *PAX6, SOX2, HES1, HES5*, and *CDH2* (**Figure 3B; Figure S2**). This is the most inclusive and heterogeneous class of cells, showing the potential of organoid-derived progenitor cells to differentiate into the majority of cell types present in the human brain. However, their abundance also highlights the early stage of development modeled in this system.

The third cell class represents the Neurons (n=4152 total cells) and is defined by expression of neuronal marker STMN2^72^ (**Figure 3C-D; Figure S2**). It includes the most immature and heterogeneous immature Neurons (iN) which express *MAP2, DCX*, some dorsal markers like NEUROD1 and NEUROD4 as well as ventral marker *OTX2*, but also some retinal markers like CRX and RCVRN. In spite of some ventral markers present, our organoids primarily seem to differentiate along a dorsal lineage, with the immature Dorsal Neurons (iDN) expressing *MAP2, DCX*, and some dorsal-specific NEUROD6 and the most differentiated Dorsal Neurons (DN) expressing NEUROD6 abundantly, as well as glutamatergic receptor *GRIA2* and synaptic protein *SNAP25* (**Figure 3B; Figure S2**).

#### Effect sizes of GR-activation response at the single-cell level

Overall, GR-activation effect sizes were moderate, consistent with the single-cell technique, with absolute fold-change rarely exceeding 2.0-fold, and a trend for larger FC in downregulated transcripts. Genes associated with neuronal maturity like *NEUROD6, C1orf61, NFIA*, and *NFIB*, were among the most strongly down-regulated in multiple clusters (**Figure 4E** inset table for FC). The DN cluster exhibited the largest FCs, but even this only had the top 6 ranked transcripts by p-value with absolute FC above 2.0 (**Figure S5, Table S8**). Interestingly, all these 6 transcripts were down-regulated by Dex, and included the neuron-specific genes mentioned above, pointing to a disruptive effect of GR-activation on neuronal differentiation and maturation.

#### Gene Ontology analyses following differential expression in single-cell transcriptome data

To understand cell-type-specific response patterns, we performed a gene ontology enrichment analysis in the DE gene lists for each of the three cell classes (**Table S9**). We focused on the Biological Process (BP) GO term class, as it was most relevant for the questions in this study. The trends identified reflected the fact that glucocorticoids in very early neurodevelopment cell types like the Neuroepithelia can impact early development-relevant biological processes, with significant terms referring to embryonic placenta development and proliferation. In the maturing cells of the Progenitors group, terms like cell differentiation, head development, and neurogenesis were enriched, reflecting a neuronal specification and consequently the effect of glucocorticoids on these developmental processes. Finally, in the Neurons cell class, the differentially-expressed genes were additionally enriched for more specialized GO terms like forebrain development and neuron development, in line with a glucocorticoid impact on later stages of neuronal and brain maturation (**Table S9**).

#### Cell-type specific glucocorticoid response in organoids is GR-dependent

Since glucocorticoid-response has previously been established by our group and others in different tissue types like peripheral blood cells^75^ as well as more recently neuronal cell types like hippocampal progenitors^24^, we wanted to test the overlap of these previous findings with our results from GC-exposure in organoids. We tested whether the effects we observed following glucocorticoid exposure was dependent on expression of the glucocorticoid receptor (NR3C1 gene). We found that cells with sufficient mRNA levels of *NR3C1* and thus positive for this gene’s expression in our data, tended to have significantly (p-value =0.01) higher absolute fold changes in all three cell classes (Neuroepithelia p-value <0.0001, Progenitors p-value = 0.01, Neurons p-value = 0.018) compared to *NR3C1*-negative cells after Dexamethasone treatment (**Figure S4**). This fold-change distribution was tested in a set of genes that was previously determined to be indicative of glucocorticoid-response in hippocampal progenitor cells following a similar acute stimulation paradigm^24^. This suggests that the Dex-response phenotype of individual cells is in part dependent on presence of the receptor.

### Supplementary Figures

**Supplementary Figure 1:**
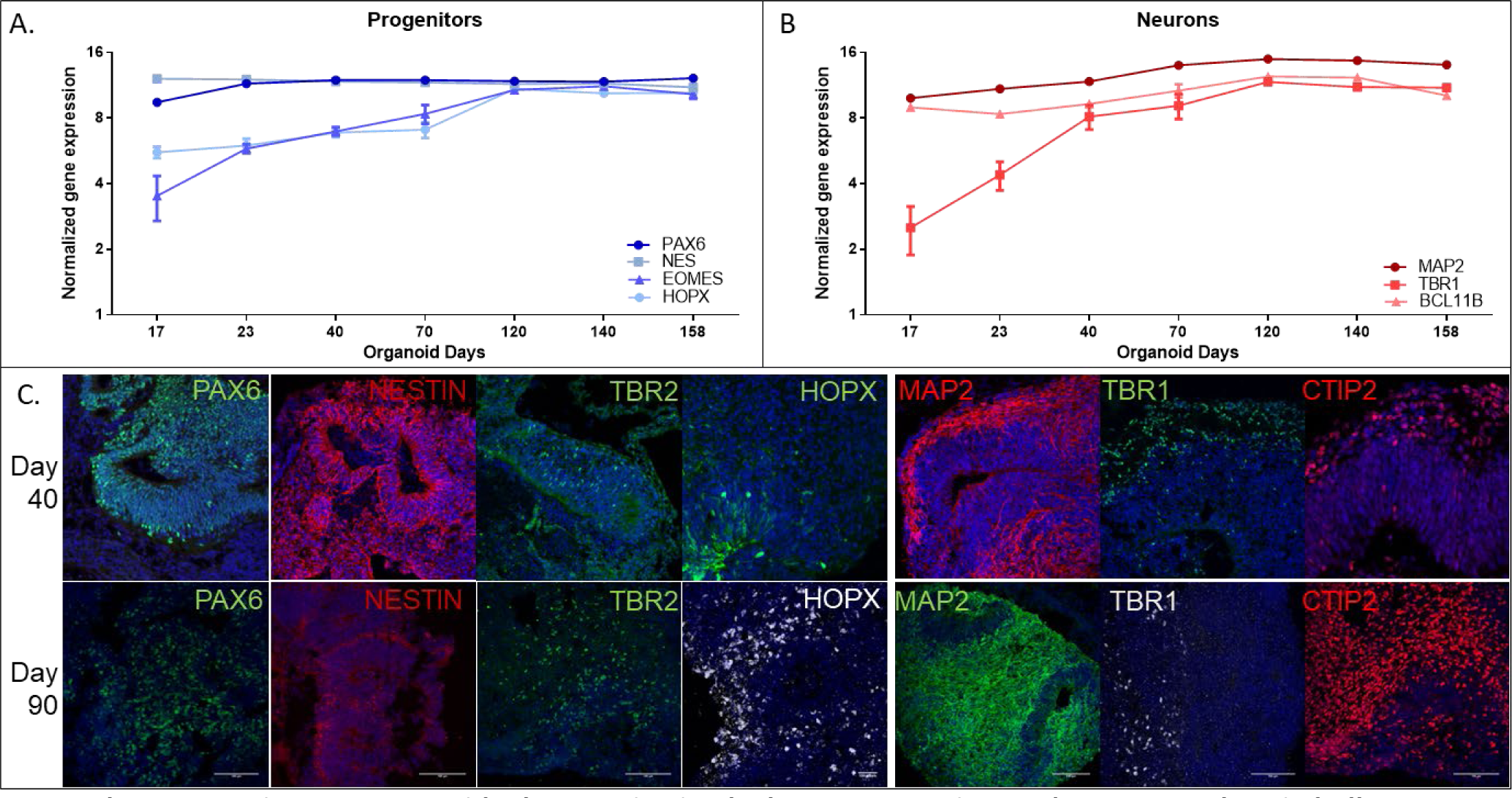
Organoids characterization by known progenitor and neuron markers in bulk RNAseq. **A-B.** At RNA expression levels in bulk RNAseq data at days 17-158 in culture. Both progenitor (blue, **A**) and neuronal (red, **B**) marker genes are chose. **C.** Using Immunofluorescence and confocal microscopy at protein expression levels at days 40 (top) or 90 (bottom) in culture. Progenitor markers: PAX6 (green), NESTIN (*NES* gene, red), TBR2 (*EOMES* gene, green), HOPX (green in day 40 and grey in day 90). Neuron markers: MAP2 (red in day 40 and green in day 90), TBR1 (green in day 40 and grey in day 90), CTIP2 (*BCL11B* gene, red). For all images in **C-D**, DAPI is used as nuclear marker (blue).

**Supplementary Figure 2:**
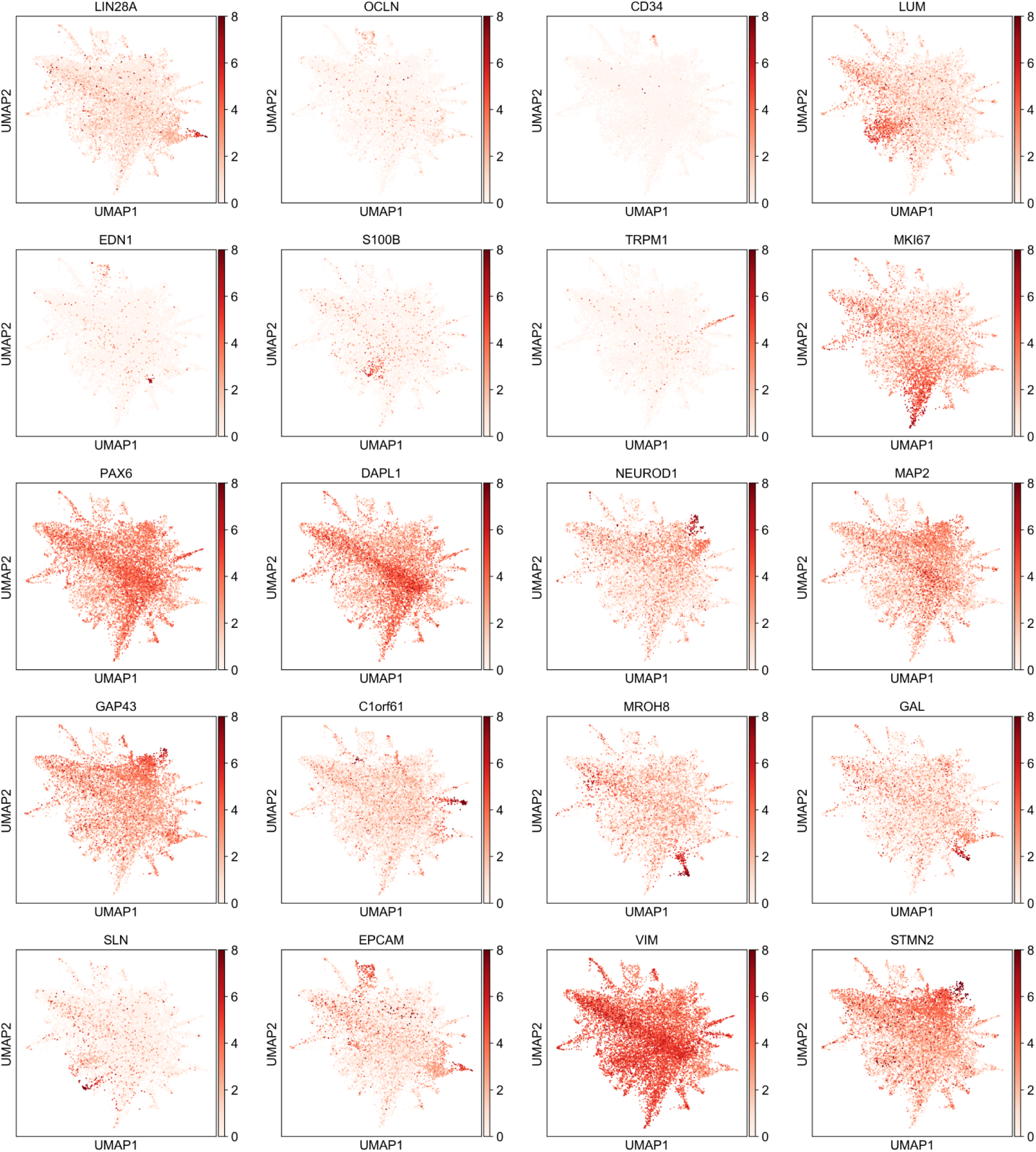
UMAPs for cell-types or cell classes. *LIN28A, OCLN, CD34, LUM, EDN1, S100B, TRPM1, MKI67, PAX6, DAPL1, NEUROD1, MAP2, GAP43, C1orf61, MROH8, GAL, SLN, EPCAM, VIM, STMN2* are referenced in dot plot in **Figure 3B** for cell-type characterization and in **Figure 3D** for cell class characterization. The last three plots represent the three cell classes defined in **Figure 3D**, as follows: Neuroepithelial cells = *EPCAM*, Progenitors = *VIM*, Neurons = *STMN2*. Detailed explanation **in Supplementary Results**.

**Supplementary Figure 3:**
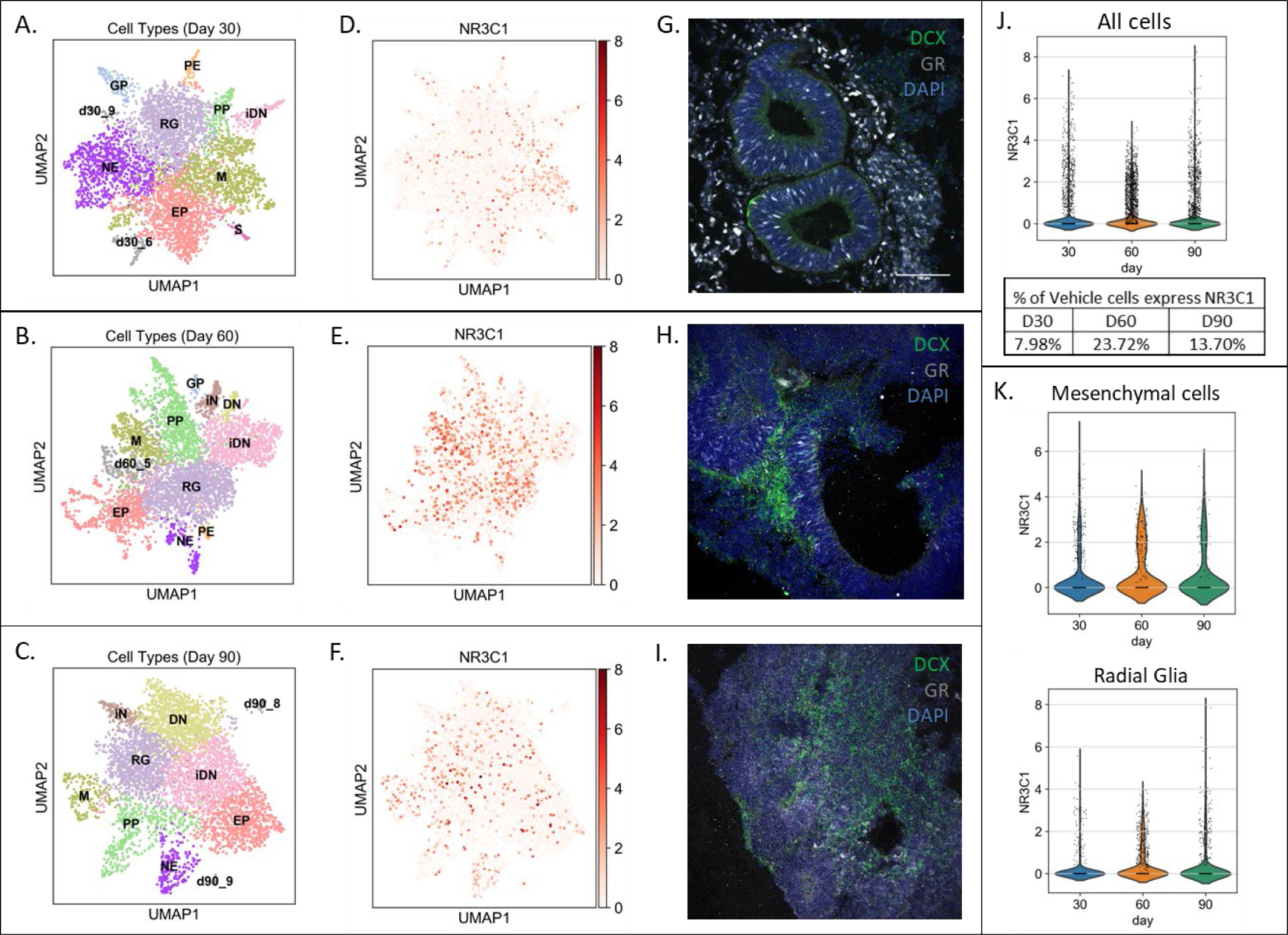
Cell type characterization by individual days. **A-C.** UMAP plot of cell-type clustering individually reported for day 30, 60, or 90. **D-F.** UMAP plot of *NR3C1* gene (GR protein) expression individually reported for day 30, 60, or 90. **G-I.** Immunofluorescence image of GR protein staining in wild-type organoids at days 30, 60, or 90. Magnification is 25x, obtained with a confocal microscope (nuclei-DAPI = blue; DCX-neurons = green; GR = white). **J.** Violin plot of *NR3C1* expression individually reported for day 30, 60, or 90 in all combined cells. Inset table: percent of total cells at each day expressing *NR3C1* at detectable levels. **K.** Violin plots of *NR3C1* expression individually reported for day 30, 60, or 90 in individual cell types of interest: left, Mesenchymal cells (M) clusters; right: Radial Glia (RG) cells clusters.

**Supplementary Figure 4:**
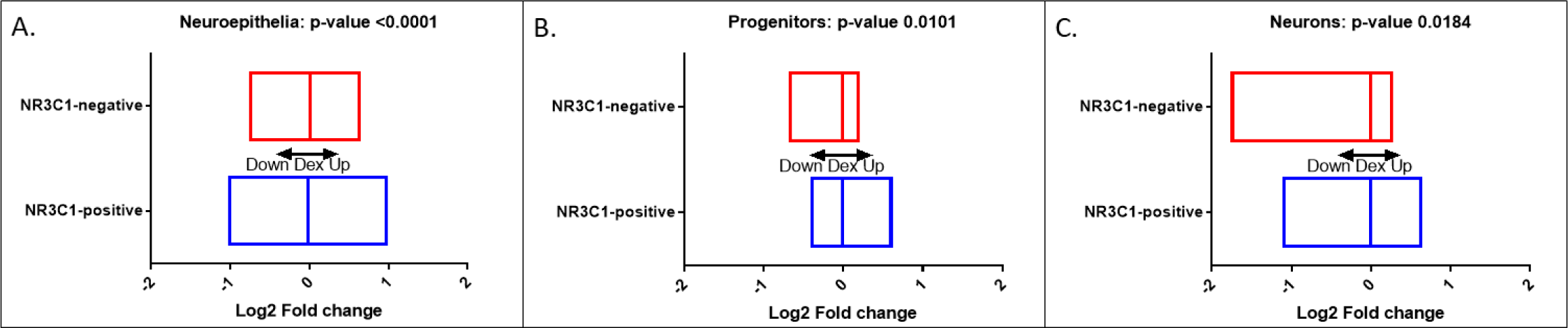
Differential gene expression by cell class in GR-positive vs. GR-negative cells. Distribution and statistics shown in **A**. for Neuroepithelial cells; in **B**. for Progenitor cells; and in **C.** for Neurons.

**Supplementary Figure 5:**
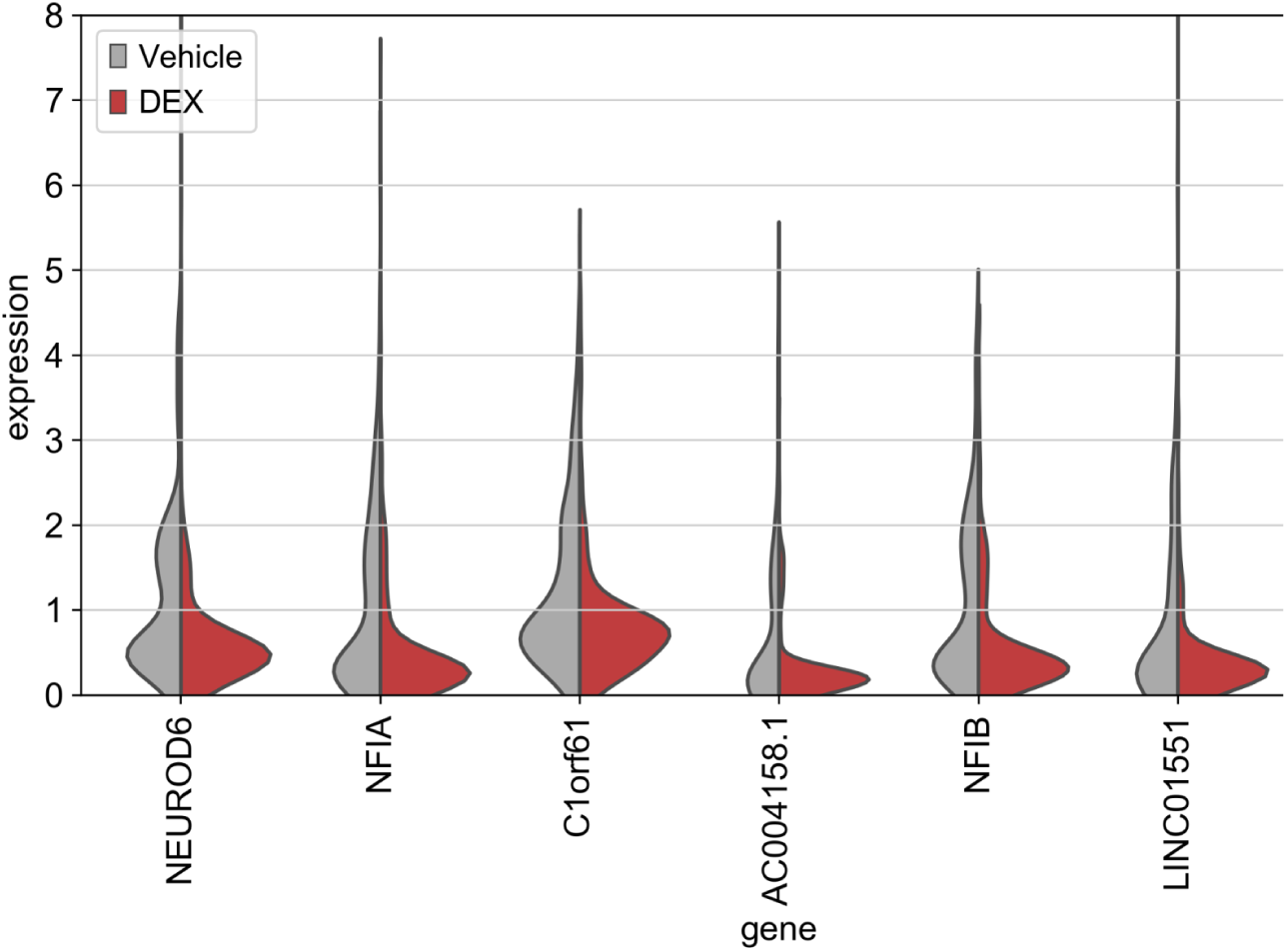
Violin plots of the top 6 DE genes from Dorsal Neurons (DN) cluster analysis in all combined cells. These genes (*NEUROD6, NFIA, C1orf61, AC004158.1, NFIB, LINC01551*) resulted in the highest (negative) fold changes across all analyses.

**Supplementary Figure 6:**
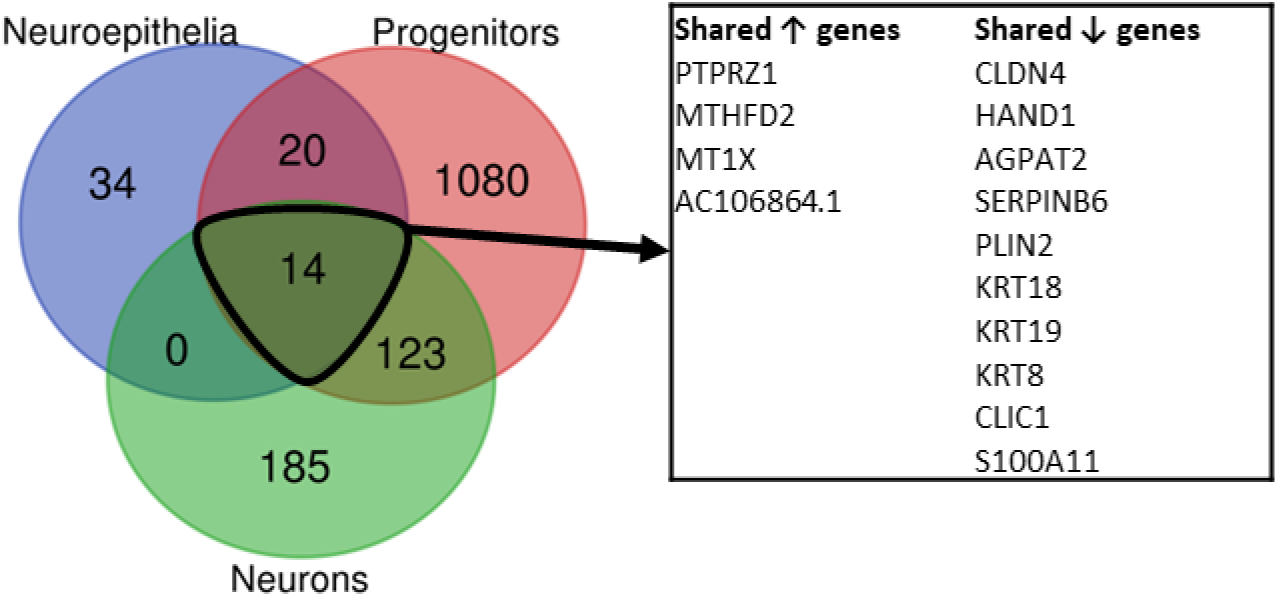
Venn diagram of overlapping differentially-expressed genes for the cell class analyses. The q-value cut-off was 0.05 for DE gene lists, and cell classes are represented as follows: Neuroepithelial cells (blue), Progenitors (red), Neurons (green).

### Supplementary Table Legends

**Supplementary Table 1:** Bulk RNAseq counts matrix. Batch-corrected and normalized gene expression counts for each of 21 RNAseq libraries, following ImpulseDE2 analysis. Genes in Column 1 are presented as Ensembl IDs. Each library/sample name is presented as “DayX_replicate”.

**Supplementary Table 2:** ImpulseDE2 results depicting genes that peak (significantly different from the other days) at each organoid developmental time point. Genes’ normalized expression levels are represented visually in **Figure 2A** via a heatmap.

**Supplementary Table 3**: Quantitative PCR differential expression after dose-time experiment in bulk organoids at day 45 in culture. Treatment was done in the medium with Dexamethasone dissolved in DMSO (10nM, 100nM, and 1000nM concentrations). The vehicle condition was DMSO. Triplicate biological experiments were used for the treatment, and the qPCR analyses were run in quadruplicate technical replicates. Analysis of qPCR data was done using the absolute quantification method.

**Supplementary Table 4:** Cluster markers.

**Supplementary Table 5:** Significant DE results for Day30 organoids, based on cluster classification (in 13 named cell-type clusters) and cell class (in 3 cells classes).

**Supplementary Table 6:** Significant DE results for Day60 organoids, based on cluster classification (in 13 named cell-type clusters) and cell class (in 3 cells classes).

**Supplementary Table 7:** Significant DE results for Day90 organoids, based on cluster classification (in 13 named cell-type clusters) and cell class (in 3 cells classes).

**Supplementary Table 8:** Significant differentially-expressed genes from all-cells combined analysis, based on cluster (in 9 named cell-type clusters with significant results) and cell class (in 3 cells classes) classification.

**Supplementary Table 9:** Gene ontology analysis in DE genes from three cell classes. Enrichment tested against Biological Process (BP) GO terms and multiple testing corrections performed using Benjamini-Hochberg correction against number of terms and number of test genes sets.

**Supplementary Table 10:** FUMA GWAS enrichment results. GWAS-significant traits with 5 or more overlapping genes to the test gene set (DE after Dex in Progenitors of Neurons in organoids) were included in the enrichment analysis. Benjamini-Hochberg multiple testing correction was done for each test gene set against the number of traits included in analysis.

**Supplementary Table 11:** Enrichment analyses of psychiatric disease-related gene sets associated genome-wide significant common variants from the genome-wide meta-analysis of the Cross-Disorder Group of the Psychiatric Genomics Consortium.

**Supplementary Table 12:** Enrichment analyses of neurodevelopmental disease-related gene sets from the DisGeNET database.

**Supplementary Table 13**: Enrichment of Genes with Loss-of-Function mutation associated with Intellectual Disability from the Developmental Brain Disorders Database (DBDD) genes database (last updated Mar 2018).

